# Mycobacteria-specific CD4^+^IFN_-γ_^+^ cell expresses naïve-surface markers and confers superior protection against tuberculosis infection compared to central and effector memory CD4+ T cell subsets

**DOI:** 10.1101/384784

**Authors:** Jinyun Yuan, Janice Tenant, Thomas Pacatte, Christopher Eickhoff, Azra Blazevic, Daniel F. Hoft, Soumya Chatterjee

## Abstract

Failure of the most recent tuberculosis (TB) vaccine trial to boost BCG mediated anti-TB immunity despite highly durable Th1-specific central (T_CM_) and effector (T_EM_) memory cell responses, highlights the importance of identifying optimal T cell targets for protective vaccines. Here we describe a novel, *Mycobacterium tuberculosis* (Mtb)-specific IFN-γ^+^CD4^+^ T cell population expressing surface markers characteristic of naïve T cells (T_NLM_), that were induced in both human (CD45RA^+^CCR7^+^CD27^+^CD95^-^) and murine (CD62L^+^CD44^-^Sca-1^+^CD122^-^) systems in response to mycobacteria. In BCG vaccinated subjects and those with latent TB infection, T_NLM_ cells, compared to bonafide naïve CD4^+^ T cells were identified by absence of CD95 expression and had increased expression CCR7 and CD27, the activation markers T-bet, CD69 and PD-1 and the survival marker CD74. Increased T_NLM_ frequencies were noted in the lung and spleen of wild type C57BL6 mice at 2 weeks after infection with Mtb, and progressively decreased at later time points, a pattern not seen in TNF-α^+^CD4^+^ T cells expressing naïve cell surface markers. Importantly, adoptive transfer of highly purified T_NLM_ from vaccinated ESAT-6_1-20_-specific TCR transgenic mice conferred superior protection against Mtb infection in Rag-/- mice when compared with total meory populations (central and effector memory cells). Thus, T_NLM_ cells may represent a memory T cell population that if optimally targeted may significantly improve future TB vaccine responses.

## Introduction

Tuberculosis (TB) disease caused by *Mycobacterium tuberculosis* (Mtb) affects more than 10 million and claims 1.5 million lives worldwide every year [1]. Mtb primarily infects lungs via the aerosol route and is the leading cause of death from a single infectious agent, ranking above HIV/AIDS. Bacillus Calmette–Guérin (BCG)-the only approved TB vaccine, is given in childhood but is poorly protective against adult onset TB [2, 3]. Studies in mouse models [4] and HIV patients [5] have shown that a lack of CD4+ T cells leads to increased disease susceptibility. A failure of disease control is also seen with a genetic deficiency of IFN-γ and IL-12 signaling [6]. Furthermore, mice specifically deficient in CD4^+^ cells producing IFN-γ (CD4^+^IFN-γ^+^) are more susceptible to TB disease [7, 8]. Although studies have shown that CD4^+^IFN-γ^+^ independent mechanisms are also involved in protection [9, 10], development of an improved TB vaccine requires better understanding of the nature of an optimal CD4^+^ and more specifically, CD4^+^IFN-γ^+^ response against Mtb. Furthermore, since recall or memory CD4^+^ responses are necessary for vaccine efficacy, it is important to understand which Mtb-specific CD4^+^ T cell memory subset provides optimal anti-Mtb immunity. BCG generates robust CD4^+^ central (T_CM_) and (T_EM_) responses but fails to provide lasting immunity [11]. Additionally, the most recent TB vaccine (MVA85) trial that failed to improve TB-specific immunity despite induction of robust T_CM_ +T_EM_ responses [12, 13]. These findings beg the question—is there a different population of CD4^+^ memory T cells that we should be targeting. Phenotypic, functional, and gene expression properties of these T-cell subsets suggest [14] that human memory T cell differentiation follows a linear progression. The continuum ranges from naïve (T_N_) to more differentiated memory subsets: central memory (T_CM_), then effector memory (T_EM_), and terminal effector (T_TE_) cells. Less-differentiated cells give rise to more differentiated progeny in response to frequency and intensity of antigen stimulation. With increasing differentiation, memory T cells progressively acquire or lose specific functions. An emerging theme is that with increasing differentiation, higher effector function is gained at the price of shorter survival and proliferative capacity. It is thus reasonable to posit that a vaccine that generates a higher frequency of less differentiated memory populations might be more effective against Mtb re-challenge. It was recently shown that a memory T cell expresses surface markers characteristic of naïve T cells (CD45RA^+^CCR7^+^) demonstrates enhanced survival and self-renewal capacity (T_SCM_) [15, 16]. These cells have been detected in BCG vaccinated infected subjects [17]. We had previously identified Mtb antigen-specific cytokine production in a subset of CD95- early precursor cells with a naïve-like phenotype [18]. Here we describe a population of Mtb-antigen specific IFN-γ^+^CD4^+^ cells with a naïve phenotype (T_NLM_) that were induced in response to mycobacterial antigens and confer superior protection against Mtb infection.

## Results

### T_NLM_ cells produce cytokines in response to mycobacterial antigens

To understand the profile of Mtb-specific memory CD4 T-cell responses, we cultured PBMCs from subjects with latent TB infection (LTBI) and Mtb unexposed (quantiFERON-TB Gold negative) healthy controls (HC). Cells were stimulated with Mtb antigens ESAT-6 or CFP-10 overnight and the frequency of different memory subsets producing cytokines IFN-γ and TNF-α was assessed by multi-parametric flow cytometry. The gating strategy shown in Fig. S1. We defined different memory subsets by combination of surface markers CD45RA, CCR7, CD27, CD95: naïve T (T_N_) cells as CD45RA^+^CCR7^+^CD27^+^CD95^-^; T memory stem cells (T_SCM_) as CD45RA^+^CCR7^+^CD27^+^CD95^+^; central memory T cells (T_CM_) as CD45RA^-^CCR7^+^; effector memory T (T_EM_) cells as CD45RA^-^CCR7^-^; terminal effector T cells (T_EMRA_) as CD45RA^+^CCR7. In the LTBI group, increased frequencies of Mtb Ag (ESAT-6 and CFP-10) IFN-γ^+^ frequencies were noted among overall CD4^+^ cells, T_CM_ and T_EM_ (expressed as net frequency to baseline), compared to HC (Figure 1a). In contrast, subjects with LTBI demonstrated higher TNF-α^+^ frequency within overall CD4^+^ and T_EM_ cells but not T_CM_ in response to ESAT-6 and CFP-10 (Figure 1a).

**Fig.1.**
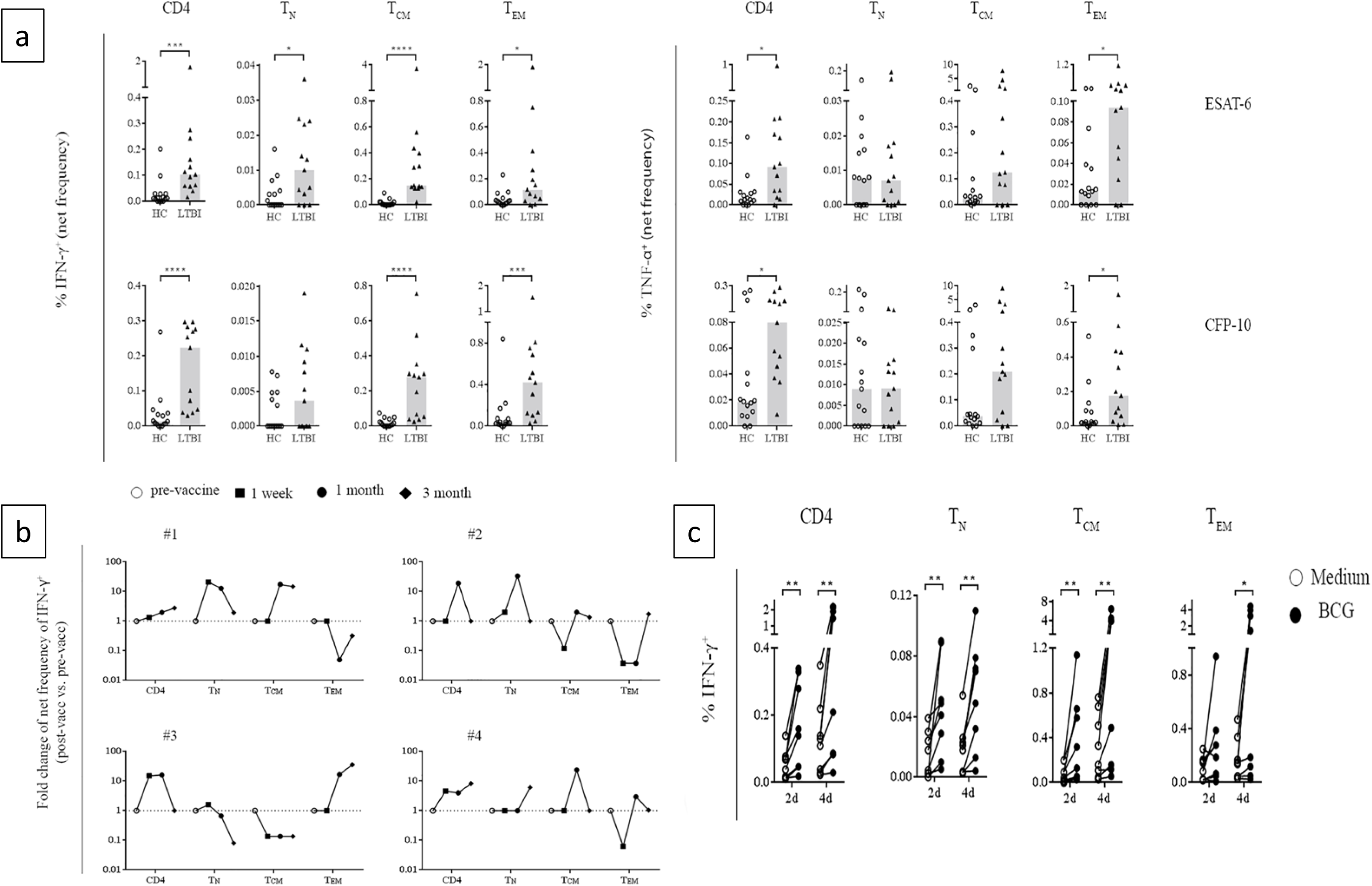
IFN-γ^+^CD4^+^ T_N_ (T_NLM_) are induced in response to mycobacterial antigens. (**a**) Frequency of IFN-γ^+^ and TNF-α^+^ cells on different CD4 T cell memory subsets in PBMCs from human subjects with latent TB infection (LTBI) and healthy controls (HC) after stimulated overnight with 10μg/mL CFP-10 or ESAT-6. Data are presented net frequencies of IFN-γ-producing CD4^+^ T cells from antigens stimulation culture compared with medium alone culture. Bar height represents median. P values were calculated using the Mann-Whitney test (*p<0.05, **p<0.01, ***p<0.001, ****p<0.0001). (**b**) Fold change of Mtb peptides-induced frequencies of IFN-γ- producing CD4^+^ T cells comparing post-vaccination (1week, 1month, 3month) with pre-vaccination in PBMCs from PPD negative and QuantiFERON TB Gold negative human subjects (V#1,2,3,4) vaccinated with BCG after stimulated overnight with Mtb-specific peptide pools. (**c**) Frequency of IFN-γ^+^ cells on different CD4 T cell memory subsets in PBMCs from HC after cultured with BCG or medium for 2d or4d following re-stimulated overnight with BCG. P values were calculated using the Wilcoxon test (*p<0.05, **p<0.01, ***p<0.001, ****p<0.0001).

Interestingly, in addition to known memory subsets, a significantly increased frequency of IFN-γ^+^CD4^+^ was observed in subjects with LTBI within cells expressing naïve-surface markers (CD45RA^+^CCR7^+^CD27^+^) in response to ESAT-6 (Median net frequency [Fo] =0.01 vs 0, p<0.05) and CFP-10 (Fo =0.00363 vs 0, p=0.056), when compared with HC. However, no such differences were noted in these cells for TNF-α production in response to ESAT-6 (Fo =0.007 vs 0.00773) and CFP-10 (Fo =0.00908 vs 0.00896). These results suggested an IFN-γ^+^ CD4^+^ “memory response” in cells expressing naïve surface markers. These cells were primarily CD95^-^ and their “memory-like” behavior along with the expression of markers of naïve T cells, led us to call these IFN-γ^+^CD4^+^ cells ‘naive like memory’ (T_NLM_).

To further examine Mtb antigen-driven recall responses within T_NLM_ cells, human PBMCs collected from BCG vaccinated subjects pre-vaccination and at 1 week, 1 month and 3 months post vaccination, were cultured with media alone or stimulated with a Mtb-specific peptide pool (kind gift from Dr. Cecilia Lindestam Arlehamn, La Jolla Institute) overnight. This panel of Mtb antigens was previously shown to elicit similar frequencies of Mtb-specific CD4 T cell responses across diverse populations [19]. Data (Figure 1c) are presented as fold changes of Mtb peptide-induced frequencies of IFN-γ-producing CD4^+^ T cells comparing post-vaccination with pre-vaccination responses. Frequency of Mtb peptide induced total IFNγ^+^CD4^+^ cells increased overtime after BCG vaccine in all 4 individuals. Moreover, increased T_NLM_ frequencies were seen after BCG vaccination in response to Mtb peptides in 3 out of 4 volunteers.

Next, to determine if similar T_NLM_ response could be elicited in vitro, PBMCs from healthy PPD-/IGRA-HC were cultured with BCG for 2 or 4 days and then restimulated overnight with BCG. On BCG restimulation of PBMCs cultured with BCG, increased frequencies of T_NLM_ were seen in compared with BCG restimulation of cells that were medium rested. This was noted in cells cultured for 2 days (Fo= 0.01126 vs 0.0450, p<0.01) or 4d (Fo= 0.0195 vs 0.0595, p<0.01). A similar trend was noted in frequencies of overall IFN-γ^+^CD4+, IFN-γ^+^T_CM_ and IFN-γ^+^T_EM_ cells (Figure 1d). This suggested that BCG induced T_NLM_ cells exhibit BCG-specific recall similar to IFN-γ^+^T_CM_ and IFN-γ^+^T_EM_ cells.

### T_NLM_ cells have a unique activated phenotype

To further characterize antigen-specific T_NLM_, we compared the expression of different markers of T cell activation and survival on IFN-γ^-^ vs. IFN-γ^+^ CD4+ memory T cell subsets. As shown in Figure 2, we compared IFN-γ^-^ naïve CD4+ T cells (T_N_) with T_NLM_, IFN-γ^-^ central memory T cells (T_CM_) with IFN-γ^+^T_CM_ (T_CM_-γ), IFN-γ- effector memory cells (T_EM)_ with IFN-γ^+^T_EM_ (T_EM_-γ) in subjects with LTBI (Figure 2) after overnight stimulation with ESAT-6. Expression levels of the activation markers CD69, CD25, FoxP3, PD-1, T-bet, survival markers CD74 and BCL-2 and phenotypic markers CCR7 and CD27 are depicted as geometric mean fluorescence intensity (gMFI). In addition, because a CD8+CD49d+ IFN-γ^+^ memory T cell with naïve phenotype was recently described to increase with age, with the frequency of these cells correlating inversely with the residual capacity of the immune system to respond to new infections with age[20], we also measured CD49d expression on the various memory subsets.

**Fig. 2.**
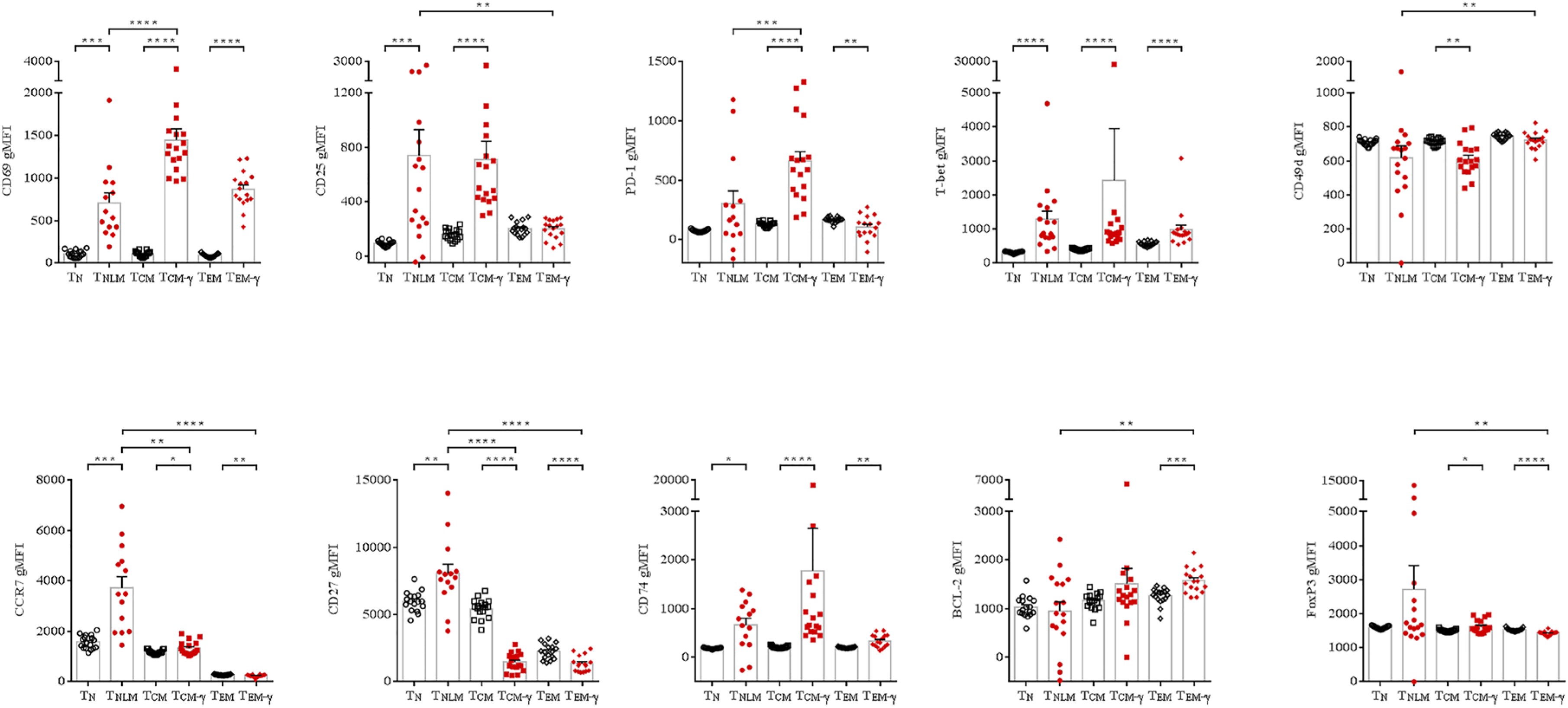
T_NLM_ exhibit distinct phenotypic and activation markers compared to T_N_, T_CM_ and T_EM._ Expression levels demonstrated as geometric mean fluorescence intensity (gMFI) of different markers involved in T cell activation and function on T_N_, T_NLM_, T_CM_, IFN-γ^+^T_CM_ (T_CM_-γ), T_EM_, and IFN-γ^+^T_EM_ (T_CM_-γ) in PBMCs from LTBI after overnight stimulated with EAT-6. Horizontal lines indicate the mean (+SEM). P values were calculated using the Wilcoxon test between T_N_ and T_NLM_, T_EM_ and T_EM_- γ, T_CM_ and T_CM_- γ and the Mann-Whitney test between T_NLM_ and T_EM_-γ, TNLM and T_CM_-γ (*p<0.05, **p<0.01, ***p<0.001, ****p<0.0001). Dunn’s correction was used for multiple comparisons.

Upon ESAT-6 stimulation, T_NLM_ compared to T_N_, showed higher expression of CD69, CD25, PD1, T-bet, CCR7, CD27, CD25, CD74 and FoxP3, negligible expression of CD95 and no difference in expression of CD49d or Bcl-2 (Figure 2). We also compared expression of these markers between T_NLM_, T_CM_-γ and T_EM_-γ. Compared to T_CM_-γ cells, T_NLM_ had lower expression of CD69 and PD1. Interestingly, T_NLM_ showed higher expression of CD25, and FoxP3 and lower expression of CD49d and Bcl-2 compared to T_EM_-γ. This suggested that in patients with LTBI, IFN-γ^+^CD4^+^ T_N_ (T_NLM_) cells expressed unique surface and activation markers in response to ESAT-6 compared to T_N,_ T_CM_-γ and T_EM_-γ.

### In vivo, Mtb-specific T_NLM_ cells are CD62L^+^CD44^-^Sca-1^+^CD22^-^ and are lost rapidly in the lung during Mtb infection

We wanted to further ascertain the kinetics of Mtb antigen-specific frequencies of T_NLM_ cells during in vivo Mtb infection. C57BL/6 (B6) mice were aerosol challenged with 100 CFU of Mtb Erdman with Glas-Col inhalation exposure systems and lung, mediastinal lymphocyte node and spleen were harvested after 4wks. As expected, ESAT-6 tetramer positive CD4 T cells were detected in Mtb infected mice with higher frequencies of ESAT-6 Tet^+^CD4^+^ presented in lung than that in spleen. To further analyze the percentage of memory subsets within ESAT-6^+^Tet^+^ CD4 T cells, CD62L and CD44 were used to identify T_N_ (CD62L^+^CD44^-^), T_CM_ (CD62L^+^CD44^+^), T_EM_ (CD62L^-^CD44^+^) and T_EFF_ (CD62L^-^CD44^-^) and Sca-1 and CD122 to identify T_SCM_ (CD62L^+^CD44^-^Sca-1^+^CD122^+^) (Fig. S2). Interestingly, the predominant ESAT-6 Tet^+^CD4^+^ cells noted within the CD62L^+^CD44^-^ population were cells expressing Sca-1^+^CD22^-^ (Figure 3a). These cells were also the predominant IFN-γ^+^ population. In this model, therefore, we defined T_NLM_ as IFN-γ^+^ CD62L^+^CD44^-^ Sca-1^+^CD22^-^.

**Fig. 3.**
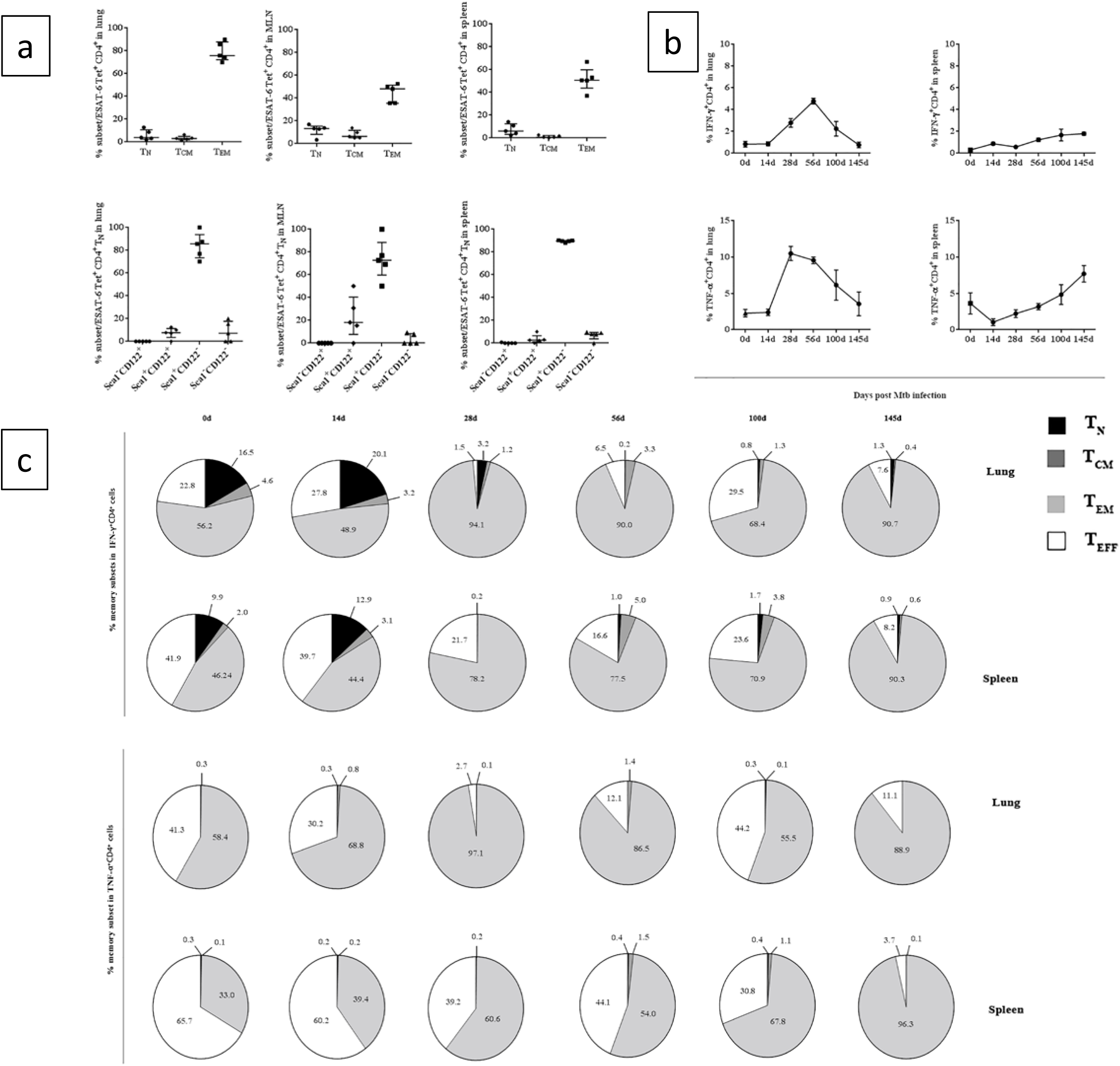
ESAT-6-specific T_NLM_ are present prior to and at early stages of Mtb infection. (**a**) Memory phenotype of ESAT-6 tetramer^+^ CD4^+^ T cells from Mtb infected B6 mice. (**b**) Frequency of ESAT-6-specific IFN-γ^+^ and TNF-α^+^ CD4^+^ T cells in lungs and spleens at different time points after Mtb infection in ESAT-6 TCR transgenic mice after re-stimulation with 10μg/mL ESAT-6_1-20_ for 6 hours ex vivo. **(c**) The memory subsets of ESAT-6-specific IFN-γ^+^ and TNF-α^+^ CD4^+^ T cells present in the lung and the spleen at the various time points shown in (b).

To further study the kinetics of Mtb-antigen-specific T_NLM_ during Mtb infection, the recall response of different memory ESAT-6-specific CD4+ subsets to ESAT-6_1-20_ was assessed from Mtb infected ESAT-6 TCR transgenic mice at various time points up to 145 days after Mtb infection. In this mouse model, Mtb growth kinetics in both lung and spleen is similar to that in wild type B6 after aerosol challenge of Mtb showing progressive increase up to 2 weeks followed by gradual plateauing. Net frequencies of ESAT-6_1-20_-specific IFN-γ^+^CD4^+^ peaked at day 56-post infection (p.i.) (Figure 3b), while TNF-α^+^CD4^+^ frequencies peaked at day 28 p.i. We then studied the different memory phenotypes making up the IFN-γ^+^CD4^+^ and TNF-α^+^CD4^+^ cells at all time points prior to and after Mtb infection. As expected, low overall frequencies of ESAT-6-specific IFN-γ^+^CD4^+^ T cell frequencies were seen both in the lung and spleen prior to infection. Following Mtb infection, both IFN-γ^+^T_EM_ and TNF-α^+^T_EM_ cells increased over time However, the highest frequencies of ESAT-6-specific T_NLM_ were present at14 days p.i. in both lung and spleen with low frequencies seen at all subsequent time points (Figure 3c). This pattern of the ESAT-6-specific T_NLM_ response was not seen in TNF-α^+^ cells expressing naïve surface markers (CD62L^+^CD44^-^), suggesting that T_NLM_ cells (IFN-γ^+^ CD62L^+^CD44^-^ Sca-1^+^CD22^-^) are a Mtb antigen-specific population involved in the IFN-γ but not in the TNF-α^+^ producing CD4+ T cell response to Mtb infection.

### T_NLM_ cells confer superior protection against Mtb infection compared to total memory cells (T_CM_+T_EM_)

To determine the capability of antigen-activated T_NLM_ to inhibit Mtb growth in vitro, equivalent cell numbers of purified total CD4 and T_NLM_ ex vivo from ESAT-6 TCR transgenic mice or expanded with ESAT-6_1-20_ peptide for 7d in vitro were co-cultured with Mtb infected BMDM (bone marrow derived macrophage) 1 day earlier. Cells were harvested after 3 days of co-culture. After 3d, in BMDM cultures without CD4+ T cells, a 10-fold increase in Mtb CFU was observed. Although there was a decrease in the Mtb load on adding CD4+ cells to BMDMs (Figure 4a), we saw no difference in the degree of Mtb growth inhibition when we compared ESAT-6_1-20_ peptide expanded total CD4+ T cells to unexpanded total CD4+ T cells. However, when purified T_NLM_ were sorted form total CD4+ T cells and put in culture with BMDMs, we saw significant inhibition of mycobacterial growth by peptide-exposed T_NLM_ compared peptide unexposed T_NLM_.

**Fig. 4.**
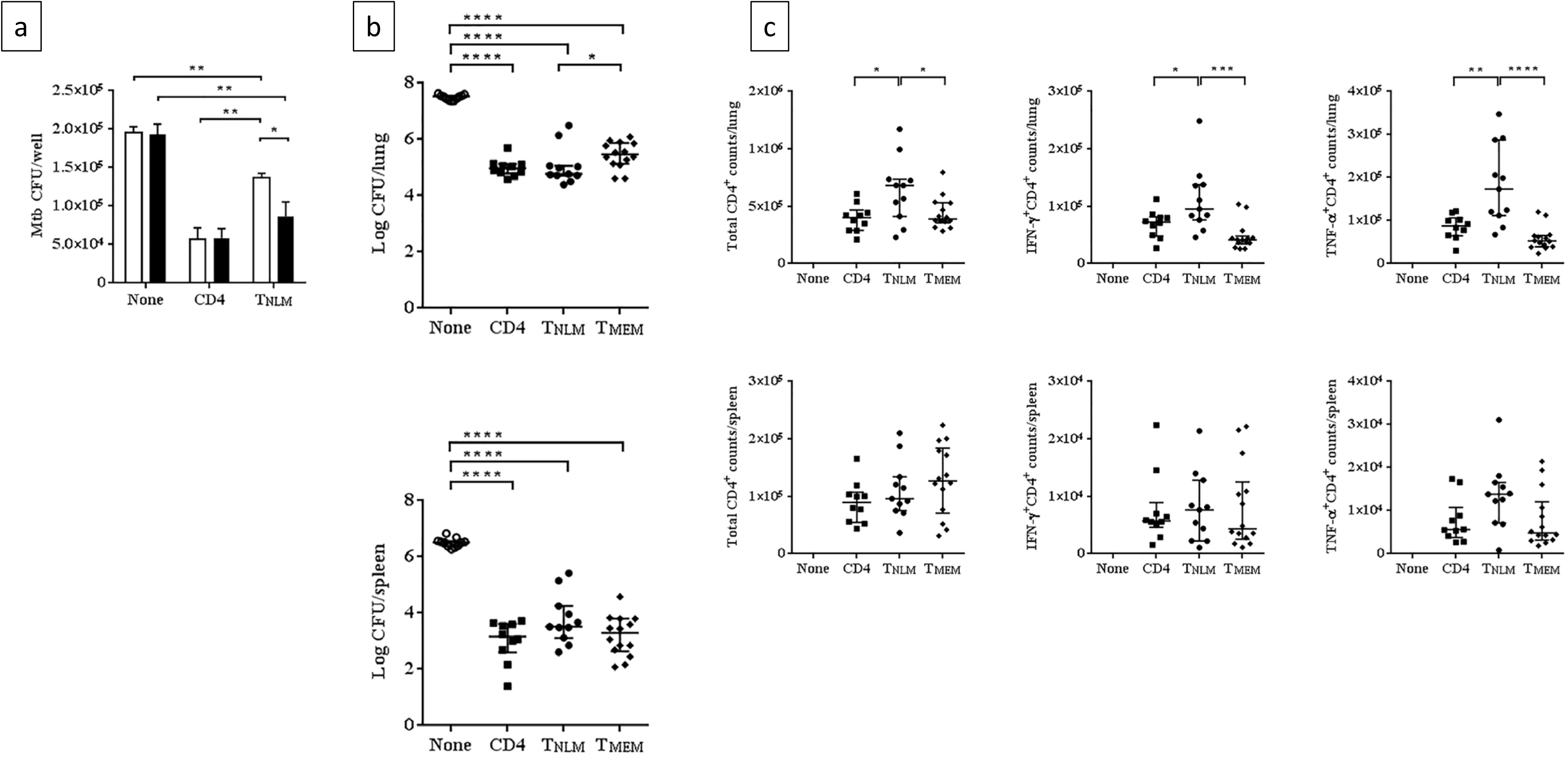
ESAT-6-specific T_NLM_ provide superior protection against TB infection in mice compared to total memory cells. (**a**) Mtb CFUs in Mtb infected BMDMs co-cultured with purified ESAT-6-specific T_NLM_ (CD4^+^CD62L^+^CD44^-^Sca1-^+^CD122^-^) and CD4 directly ex vivo or expanded with ESAT-6 _1-20_ for 7d from ESAT-6 TCR transgenic mice. (**b**) Mtb CFUs in lungs and spleens from Rag-/- mice receiving purified ESAT-6-specific CD4, T_NLM_ (CD4^+^CD62L^+^CD44^-^Sca1-^+^CD122^-^) and T_MEM_ (CD4^+^CD62L^-^CD44^+^, CD4^+^CD62L^-^CD44^-^ and CD4^+^CD62L^+^CD44^+^) 30d after Mtb infection. (**c**) Total CD4^+^, ESAT-6-specific IFN-γ^+^CD4^+^, ESAT-6-specific TNF-α^+^CD4^+^ T cell counts in lungs and spleens recovered from mice in (b). The data are representative of two independent experiments of similar design. Horizontal lines indicate the median. P values were calculated using the Mann-Whitney test (*p<0.05, **p<0.01, ***p<0.001, ****p<0.0001).

To further define whether antigen-activated T_NLM_ are important in protection from Mtb infection in vivo, equivalent cell numbers of purified total CD4, T_NLM_, and T_MEM_ (T_CM_+T_EM_) (0.5-1×10^6^) from ESAT-6 TCR transgenic mice after ESAT-6_1-20_ vaccination were adoptively transferred via tail vain injection into Rag-/- recipients who were challenged the following day with low dose (~100 cfu) Mtb. Regardless of the type of CD4 T cells transferred initially, almost all cells were found to be phenotypically T_EM_ at 28 days post-infection. However, increased total CD4+ and CD4+IFN-γ^+^ and CD4+TNF-α^+^ cells were found in lung but not in spleen from mice receiving T_NLM_ compared to those receiving total CD4+ cells and T_MEM_ (Figure 4b and 4c). More importantly, mice given T_NLM_ had lower bacterial burdens (Median frequency= 50878 vs 293082, p<0.05) than mice receiving T_MEM_ cells (Figure 4b). These results provide direct support for superior protection provided by T_NLM_ compared to overall CD4 and T_MEM_.

## Materials and methods

### Human subjects

All individuals were examined and samples collected as part of registered protocols approved by the Institutional Review Board of Saint Louis University (SLU) School of Medicine (Protocol #22975 and 26527).

4 PPD negative and QuantiFERON TB Gold negative subjects were vaccinated with BCG and peripheral blood mononuclear cells (PBMCs) were collected pre-vaccination, 1 week, 1 month and 3 months post vaccination. Subjects with latent tuberculosis infection (LTBI) were defined as those with a positive Interferon Gamma Release Assay (T-spot or quantiFERON TB Gold) test, no clinical features of active TB disease and no abnormal findings on chest X-ray. PBMCs from subjects with LTBI were collected prior to starting treatment with Isoniazid (pre-treatment) and at 3months and 6 months after starting treatment.

### In vitro short stimulation of human PBMCs

Cultures on PBMCs were performed to determine memory subsets and levels of intracellular cytokines. Briefly, cells were cultured in RPMI 1640, with 10% FBS with penicillin– streptomycin (100 U per 100 mg/ml), L-glutamine (2 mM) at 1×106 cells/FACS tube in 500ul volume. PBMCs from BCG vaccinated volunteers were stimulated overnight with Mtb-specific peptide pool (obtained from Dr. Cecilia Lindestam Arlehamn, La Jolla Institute)[19] and PBMCs from LTBI and healthy controls (HC) were stimulated overnight with 10μg/mL ESAT-6, 10μg/mL CFP-10 or positive control CytoStim (Miltenyi Biotec) in presence of 1μg/mL αCD28/CD49d co-stimulatory molecules and protein transport inhibitor GolgiStop™ (BD) was added after 4hrs. PBMCs cultured with medium alone were served as unstimulated control. After stimulation, cells are harvested and stained with fluorochrome-conjugated antibodies.

### T cell phenotyping and cytokine production

Gating was performed on live single CD4^+^ T cells. Different memory subsets are defined by combination of presence and absence of surface markers CD45RA, CCR7, CD27, CD95: naïve (T_N_ CD45RA^+^CCR7^+^CD27^+^CD95^lo^); T memory stem cells (T_SCM_ CD45RA^+^CCR7^+^CD27^+^CD95^hi^); central memory (T_CM_, CD45RA^-^CCR7^+^); effector memory (T_EM_, CD45RA^-^CCR7^-^); terminal effector (T_EMRA_, CD45RA^+^CCR7^-^). Data are depicted as frequency of CD4^+^ T cells expressing cytokines IFN-γ or TNF-α. The gating strategy is presented in Supplemental Fig. 1. Baseline values following medium culture are depicted as absolute frequency, and frequencies following stimulation with antigens are depicted as net frequency (Antigen-stimulated condition − Unstimulated condition).

### Mouse strains

C57BL/6 (B6) mice were purchased from The Jackson Laboratory (Bar Harbor, ME). ESAT-6 TCR transgenic (Tg) mice expressing the αβTCR specific for the IAb-presented ESAT-6 _1-20_ peptide were kindly provided by Dr. David L. Woodland (Trudeau Institute, Saranac Lake, NY). Rag1-/- mice on B6 background were kindly provided by Dr. Richard Di Paolo (Saint Louis University, St. Louis, MO). All mice were bred in house and maintained under specific pathogen-free conditions. Mtb infected mice (8–12 weeks old of sex matched) were housed at the Association for the Assessment and Accreditation of Laboratory Animal Care-approved BSL3 facility at the SLU per the National Research Council Guide for the Care and Use of Laboratory Animals and used in accordance with protocols established by the Institutional Animal Care and Use Committee of the Department of Comparative Medicine, SLU School of Medicine. Sample sizes were based on previous experience, and sample size calculations were approved by the Saint Louis University Animal Care Committee following AAALAC guidelines and recommendations. Although no formal blinding was done, all key experimental results were reproducible in multiple experiments.

### Aerosol infections and bacterial load determination

M. tuberculosis, Erdman strain (TMCC 107) was grown from low-passage seed lots in Middlebrook liquid medium 7H9 (BD) containing 10% ADC (BD) and 0.05% Tween 80 to mid-log phase, then aliquoted and frozen at −70°C until use. Mice received aerosol infection with the Erdman strain of M. tuberculosis by using a Glas-Col aerosol generation device (Glass-Col, Terre Haute, IN) to deliver ∼100 CFU/animal. To determine the bacterial load post infection, left lung and part of spleen from Mtb infected mouse were homogenized in 7H9 media with 0.05% Tween 80. 500μL of 1/5 diluted lung and undiluted spleen homogenate were then added to a MGIT tube and incubated in a BACTEC MGIT 320 (BD Diagnostics, Sparks, MD) liquid culture system until registered positive.

To convert time to positivity (TTP) to bacterial numbers (CFU), a standard curve was used. To produce the standard curve, 500μl of 10-fold dilutions of the mycobacterial strains spiked in mouse lung homogenate or 7H9 media with 0.05% Tween 80 were inoculated into the MGIT tubes, and TTP was plotted against CFU obtained from plating aliquots of the mycobacteria on 7H11 agar plates containing 10 % OADC supplement (BD) and 0.5 % glycerol. A linear regression analysis was carried out using GraphPad Prism version 6, and the resulting equation was used to convert TTP to CFU. Data are presented here as total number of CFUs per sample, as determined by use of a standard curve (Fig. S3).

### Preparation of single-cell suspensions

Spleens were smashed using the plunger end of a 3-c.c. syringe until completely dissociated. Lungs were cut into small pieces and incubated in RPMI medium containing collagenase XI (1 mg/ml; Sigma-Aldrich) and type IV bovine pancreatic DNase (50 mg/ml; Sigma-Aldrich) during 1hr at 37°C. The digested lungs were disrupted by gently pushing the tissue through a 40μM cell strainer. The lung and spleen single-cell suspension was lysed with red blood cells and washed for staining with appropriate fluorochrome-conjugated antibodies.

### In vitro intracellular Mtb growth inhibition assay

To assess the capability of different memory subsets of ESAT-6-specifc CD4 to inhibit intracellular Mtb growth in vitro, we had adapted mycobacterial growth inhibition assay (MGIA). Details of this method have been previously reported [21]. Briefly, mononuclear cells from the bone marrow of C57BL/6 mice were plated at 5X10^5^/ml on 6-well plate in complete culture medium containing 50ng/ml M-CSF. At day 7, macrophages were harvested and seeded into 96- flat-bottom well plate at 2X10^4^/well in 100 μL culture medium before infected with 100 μL Mtb Erdman at MOI=0.5 overnight. During the same week, splenocytes from ESAT-6 TCR transgenic mice were stimulated with ESAT-6_1-20_ 1μg/mL. At day 8, antigen expanded culture and fresh splenocytes from ESAT-6 transgenic mice were sorted out CD4^+^, T_NLM_ (CD4^+^CD62L^+^CD44^-^) by BD FACSAria IIIu. 2X10^5^ cells of each purified population were added to each well containing 2 × 10^4^ Mtb infected bone marrow derived macrophages (BMDM) (effector: target=10:1). At day 4, culture supernatants were carefully removed without disturbing the bottom cell layer and 100 μL 0.2% saponin were added to each well for 1hr at 37°C to lyse cells and release Mtb. Each cell lysate100 μl was then added to a MGIT tube and incubated in a BACTEC MGIT 320 liquid culture system. Bacterial cfus were determined as described above.

### Adoptive transfer

ESAT-6 Tg mice were subcutaneously vaccinated with 100μg ESAT-6_1-20_ emulsified in 250μg dimethyl dioctadecylammonium bromide (DDA) with 25μg monophosphoryl lipid A (MPL) and 25μg trehalose dicorynomycolate (TDM) three times at 2-week intervals. After 4wks, splenocytes from immunized mice were sorted out CD4^+^, T_NLM_ (CD4^+^CD62L^+^CD44^-^Sca1-+CD122^-^) and T_MEM_ (CD4^+^CD62L^-^CD44^+^, CD4^+^CD62L^-^CD44^-^ and CD4^+^CD62L^+^CD44^+^) by flow cytometry. 5×10^5^-1×10^6^ each purified cell population was adoptively transferred to Rag-/- mice by tail vein injection. Next day, all recipients were challenged with a low dose of Mtb Erdman (50-100 CFU/lung) by aerosol infection (Glas-Col). At 4wks of Mtb infection, lungs and spleens were removed from mice for bacterial cfu determination, cell culture with antigens and flow cytometry.

### Detection and recall response of Mtb-specific CD4 T cells

PE-conjugated tetramers (ESAT-6 4-17: I-Ab) were obtained from the National Institutes of Health Tetramer Core Facility. Single cell lymphocyte preparations were stimulated with 10μg/mL ESAT-6_1-20_ peptide in present of GolgiPlug (BD) at 37°C for 6hrs and were stained with the tetramers and incubated at 37°C for 1 h before surface and intracellular staining.

### Flow cytometry analysis

Single cell suspensions were stained for surface markers at 4°C for 30 mins. For intracellular staining, cells were further incubated with intracellular fixation & permeabilization buffer (eBioscience) and followed by staining intracellular proteins in permeabilization buffer (eBioscience) for 30 mins at 4°C. LIVE/DEAD™ Fixable Dead Cell Stain Kit (Life Technology) was used to determine the viability of cells prior to the fixation and permeabilization required for intracellular antibody staining. Purified anti-mouse CD16/CD32 mAb (BD Biosciences) and Human FcR Blocking Reagent (Miltenyi Biotec) were used to prevent nonspecific binding of Abs to the Fc receptors.

### Data acquisition and analysis

Data from stained cells were acquired using BD LSRFortessa X-20 and FACSDiva software (BD Biosciences) and were analyzed using FlowJo software (TreeStar). Statistical analysis and graphical representation of data were done using GraphPad Prism software. Median frequencies were used for measures of central tendency. Statistical significance was determined by Mann-Whitney or Wilcoxon test or Kruskal–Wallis test with Dunn multiple comparison test for multiple comparisons where indicated and denoted as *, P ≤ 0.05; **, P ≤ 0.01; ***, P ≤0.001; and ****, P ≤ 0.0001.

## Discussion

Our results suggest that IFN-γ^+^CD4^+^ T a memory cell with naïve phenotype (T_NLM_) was part of the CD4+ recall response to mycobacterial antigens in subjects with LTBI as well as those vaccinated with BCG. This was not seen with TNF-α^+^CD4^+^ T cells in either human subjects with LTBI or a murine model during Mtb infection, suggesting that T_NLM_ can be identified by IFN-γ production in response to Mtb. We found a unique expression profile of different phenotypic and activation markers in T_NLM_ cells in response to ESAT-6 when compared to T_N_ or to IFN-γ^+^ memory populations i.e. T_CM_-γ and T_EM_-γ. In wild type C57BL/6 mice ESAT-6 tetramer positive T_NLM_ were detected in Mtb infected mice and were CD62L^+^CD44^-^ Sca-1^+^CD22^-^. Similar to BCG vaccinated subjects and those with LTBI, T_NLM_ were an important component of the CD4+ IFN-γ^+^ lung response prior to and at 2 weeks’ post Mtb infection in an ESAT-6 TCR transgenic mouse model. Decrease in frequencies of ESAT-6-specific T_NLM_ with a concurrent increase in T_EM_ cells were seen as Mtb bacterial loads peaked and then plateaued. Finally, ESAT-6-specific T_NLM_ from vaccinated EAST-6 TCR transgenic mice, conferred superior protection against Mtb infection compared with total memory and effector T cells when adoptively transferred into Rag-/- mice.

It was recently shown that there is a subset of memory cells with enhanced survival and self-renewal capacity (T_SCM_) [22]. These cells express surface markers of naïve-cells (CD45RA^+^CCR7^+^) along with memory marker CD95. In contrast to T_EM_, T_SCM_ can reconstitute the entire human memory T cell repertoire after bone marrow transplantation [23]. Remarkably, these cells persisted at stable frequencies 25 years after yellow fever vaccination while T_EM_ and T_CM_ decreased over time [16]. Furthermore, CD45RA^+^ CCR7^+^ CD27^+^ *Mtb*-Tetramer^+^ positive cells were recently reported in human subjects with LTBI but not in healthy controls [24]. More than 50% of these tetramer positive cells were CD95^-^. In that study, CFP10-tetramer^+^ T_NLM_ cells clustered with bulk CD4^+^ T_SCM_ cells and were distinct from bulk CD4^+^ T_N_ cells. Tetramer^+^ T_NLM_ expressed significantly higher protein levels of CCR5, CCR6, CXCR3, granzyme A, granzyme K, and granulysin than bulk T_N_ cells.

We hypothesized that TB vaccines generating a high frequency of these CD4^+^ memory cells with “stem-cell” like properties might confer improved protection on Mtb challenge. Our results suggest that a CD4^+^ memory T cell population that expressed markers of naiïve cells (CD45RA^+^CCR7^+^) is part of the Mtb antigen-specific CD4+IFN-γ^+^ but not CD4+TNF-α memory response in subjects infected with Mtb or vaccinated by BCG. We and others have previously reported a mycobacteria-specific cytokine production by subset of naïve like cells behaving similar to antigen experienced cells [17, 18] as well as their long term persistence after treatment of tuberculosis [25]. In the current study, compared to the IFN-γ^-^T_N_, T_NLM_ expressed increased protein levels of markers of T cell activation CD69, PD-1 as well as T-bet. The expression level of these activation markers on T_NLM_ were similar or lower than that expressed by T_CM_-γ^+^ and T_EM_-γ^+^. In addition, T_NLM_ compared to IFN-γ^-^T_N_ had higher expression of HLA class II histocompatibility antigen gamma chain CD74 which has been implicated in memory T cell survival and homeostasis [26]. A naiïve-like, subset was also recently identified in CD8+ populations that increase with aging and respond to chronic viral infections. These cells termed CD8+ memory cells with naïve phenotype or CD8+T_MNP_, had high expression of CD49d relative to T_N_, T_CM_ and T_EM_. However, we did not detect higher expression of CD49d on T_NLM_ compared to the other subsets. These findings suggest that T_NLM_ cannot be distinguished from IFN-γ^-^T_N_ using CD49d. Interestingly, we saw higher expression of CD25 and FoxP3 expression on T_NLM_ compared to T_EM_-γ^+^. The implications of these results are unclear at this time but it has been suggested that IFN-γ plays a major role in induction of Tregs and a fraction of these Tregs differentiate into Th1 cells after resolution of the immune response[27, 28]. We utilized the ESAT-6_1-20_/I-A^b^-specific TCR transgenic mouse to understand the kinetics of antigen-specific T_NLM_. This model has been previously used to study priming and activation of naïve T cells [29]. It was shown previously demonstrated that high frequencies of CD62L^+^ CD44^+^ cells were present not only in the mediastinal lymph node and spleen but also in the lung of Mtb uninfected mice within 2 weeks after infection. In this study, we found high frequencies of T_NLM_ but not TNF-α+ CD4+ T cells expressing naïve surface markers prior at 2 weeks post infection. While T_NLM_ frequencies progressively decreased over time as Mtb loads increased and plateaued, there were negligible frequencies of these naïve-TNF-α^+^ cells throughout the course of infection further suggesting that T_NLM_ response might be IFN-γ-specific.

In a previous study bulk CD62L^+^CD44^-^ CD4^+^ T_N_ cells from BCG vaccinated mice were found to provide superior protection against Mtb infection in Rag-/- mice compared to T_EM_ [30]. However, there was no direct evidence provided in that study that an Mtb-antigen-specific memory subset exists within total T_N_ population. Since frequencies of tetramer positive T_NLM_ were significantly lower compared to T_CM_ or T_EM_ we utilized vaccinated ESAT-6 TCR transgenic mouse to adoptively transfer antigen-specific CD4+ T cell populations into Rag-/- mice. In addition, using an in vitro killing assay, we first confirmed that purified antigen exposed T_NLM_ from ESAT-6 TCR transgenic mice lowered bacterial loads more efficiently than T_NLM_ previously unexposed to antigen. The major population of T_NLM_ transferred was CD62L^+^CD44^-^ Sca-1^+^CD122^-^. In contrast to the previous study, we saw lowering of Mtb bacterial loads in the lungs but not in the spleens of Rag-/- mice given TNLM compared to those receiving ESAT-6- specific total memory population (T_CM+_ T_EM_).

Our study did not formally assess whether IFN-γ production is essential for the protective role of T_NLM_ against Mtb. This is especially relevant as it was shown recently that in mouse models IFN-γ accounts for only ~30% of CD4 T cell-dependent cumulative bacterial control in the lungs and excess IFN-γ production may exacerbate lung pathology [31]. Additionally, we assessed the kinetics of T_NLM_ specific for ESAT-6 and recent data seems to suggest that T_SCM_ specific for Mtb antigen Ag85B secrete primarily IL-2, while BCG and CFP-10 specific cells produced IFN-γ, TNF-α and IL-2 [24]. Although we found ESAT-6-specific T_NLM_ showed superior protection against Mtb compared to conventional memory subsets, it is not known what antigen-specificities of T_NLM_ need to be generated by a TB vaccine to generate optimal, durable CD4+ memory response. Finally, the low frequencies of these cells detected in the peripheral blood make it challenging to induce large number of these cells by vaccination. However, long-term proliferative potential of CD4+ T cells after BCG vaccination was correlated with frequencies of BCG-specific stem cell like memory cells [24] suggesting that frequencies of these cells generated by a TB vaccine might be utilized as a marker of durable CD4+ T cell responses.

Thus, our findings add to the evolving new paradigm of CD4+ memory T cells and have important implications for future rational vaccine design and host-directed therapy for tuberculosis.

